# The role of AP-1 in self-sufficient proliferation and migration of cancer cells and its potential impact on an autocrine/paracrine loop

**DOI:** 10.1101/271536

**Authors:** Sherif Ibrahim Abd El-Fattah, Aierken Abudu, Eugenia Jonhson, Neelum Aftab, Michele Fluck

## Abstract

The activating protein-1 (AP-1) family members are highly expressed in invasive cancers, but the consequences of this are not completely understood. The aim of this study was to explore the significance of elevated levels of AP-1 family members under conditions that restrict growth. We observed that invasive MDA-MB-231 cells express high levels of Fra-1, c-Jun and, Jun-D during serum starvation and throughout the cell cycle compared to non-tumorigenic and non-invasive cell lines. We then analyzed Fra-1 levels in additional breast and other cancer cell lines. We found a correlation between the high levels of Fra-1 during serum starvation and the ability of the cells to proliferate and migrate under these conditions. Utilizing a dominant negative construct of AP-1, we demonstrated that proliferation and migration of MDA-MB-231 in the absence of serum requires AP-1 activity. Finally, we observed that MDA-MB-231 cells secrete factors(s) that induce Fra-1 expression and migration in non-tumorigenic and non-metastatic cells and that both the expression of and response to these factors require AP-1 activity. These results suggest the presence of an autocrine/paracrine loop that maintains high Fra-1 levels in aggressive cancer cells, enhancing their proliferative and metastatic ability and affecting neighbors to alter the tumor environment.

## INTRODUCTION

Activating protein-1 (AP-1) is a dimeric transcription factor typically comprised of one member each from the Fos and Jun families [1, 2]. The Fos family includes 4 proteins (c-Fos, FosB and Fos related antigen 1 and 2 (Fra-1 and Fra-2)), while the Jun family is formed of 3 proteins (c-Jun, JunB and JunD) [2]. Fos and Jun genes and their proteins were discovered as mediators of tumor promotion. A common sequence (TGAG/CTCA) was identified upstream of the tumor promotor phorbol ester-induced genes, and AP-1 was isolated as a protein that binds to this sequence [3]. Approximately at the same time, c-Fos and c-Jun were determined to be cellular homologs of viral oncogenes [4–6]. Soon after that, they were recognized as components of AP-1 that together bind the AP-1 response sequence. [7, 8]. Early studies of Fos and Jun demonstrated their essential role in various steps of the cell cycle in response to treatment with growth factors [9, 10]. When expression levels of AP-1 family members were examined in non-tumorigenic cells during serum starvation and release, a common pattern was identified. Expression of all family members is low in serum-starved cells. After induction with a mitogen or serum, c-Fos is the first mRNA synthesized, followed by c-Jun expression, allowing the formation of c-Fos/c-Jun heterodimers [11]. c-Fos expression then decreases [10] and Fra-1 increases, resulting in a shift from c-Fos/AP-1 to Fra-1/AP-1 and presumably helping cells past the restriction point by inducing cyclin D1. This change in expression occurs via DNA binding in which c-Fos induces Fra-1 via an AP-1 binding site in the Fra-1 promoter [12] and in return Fra-1 turns off c-Fos by binding to the c-Fos promoter [13].

Fra-1 and c-Jun protein levels were reported to be significantly higher in aggressive breast cancer cell lines (e.g. MDA-MB-231) compared to non-invasive breast cancer cells types such as MDA-MB-468 and MCF7 [14–17]. High Fra-1 and c-Jun were found to play a role in the metastatic abilities of these cells through induction of genes that enhance cell migration and invasion such as MMP2 and MMP9 [17] and/or repression of genes that suppress these processes like TSCL1 [18]. Three kinase pathways were found to contribute to the high levels of Fra-1 in highly metastatic breast cancer cell lines. The first is the canonical MEK/ERK pathway that normally induces AP-1 activity under serum induction. The activity of this pathway in metastatic breast cancer cell lines occurs through increased activity of PKC θ, which also works through the JNK pathway [16]. The third pathway is the PI3K pathway, acting through AKT [15]. One of the hallmarks of cancer disease is the ability of malignant cells to persist and grow independent of extracellular regulatory molecules such as growth factors. Several mechanisms were proposed to account for this, including the ability of these cells to secrete their own autocrine growth-inducing factors [19]. Given the role of AP-1 in mediating multiple cancer related functions, we focused on its role in sustaining cell proliferation and migration *in vitro* in the absence of growth factors. Our results show that certain AP-1 family members are maintained at high levels in the absence of serum in aggressive cancer cells from different tissue origins, and that this enables cells to proliferate and migrate. Finally, we investigated the potential contribution of an autocrine/paracrine loop to this function of AP-1.

## Results

### Expression patterns of AP-1 members during the cell cycle in breast cancer cell lines

Studies of exponentially growing breast cancer cells have demonstrated that some AP-1 family members such as Fra-1 and c-Jun are highly expressed in invasive cell lines compared to less invasive ones [15, 17]. Given the role of AP-1 in the regulation of normal cells into the cell cycle, we sought to determine if the expression of different AP-1 members is deregulated during the cell cycle in more invasive cell lines. We analyzed expression of Fra-1, c-Fos, c-Jun, and Jun-D during serum starvation and re-entry into the cell cycle in a panel of cell lines representing non-tumorigenic (MCF10A), tumorigenic non-invasive (MDA-MB-468) and invasive (MDA-MB-231) breast cancer cell lines. As shown in Figure 1A, most of the AP-1 family members’ expression is low in serum starved MCF10A and MDA-MB-468 cells and increased upon serum addition, similar to what was first described in mouse fibroblasts [9, 10]. Both cFos and c-Jun increase early after introduction of serum, while Fra-1 and JunD increases later, and this is associated with a reduction in the level of c-Fos, which return to basal levels at 12 hours. In contrast, in MDA-MB-231 cells, expression of all family members is maintained during serum deprivation, and some family members (Fra-1 and JunD) are expressed at very high levels. The only exception in MDA-MB-231 is c-Fos, whose temporal pattern was similar to normal cells. Additionally, we used RT/qPCR to show that the change of patterns of expression of Fra-1, c-Jun and Jun-D occurs also on the mRNA level in MDA-MB-231 cells compared to MDA-MB-468 cells (Figure S1).

**Figure 1:**
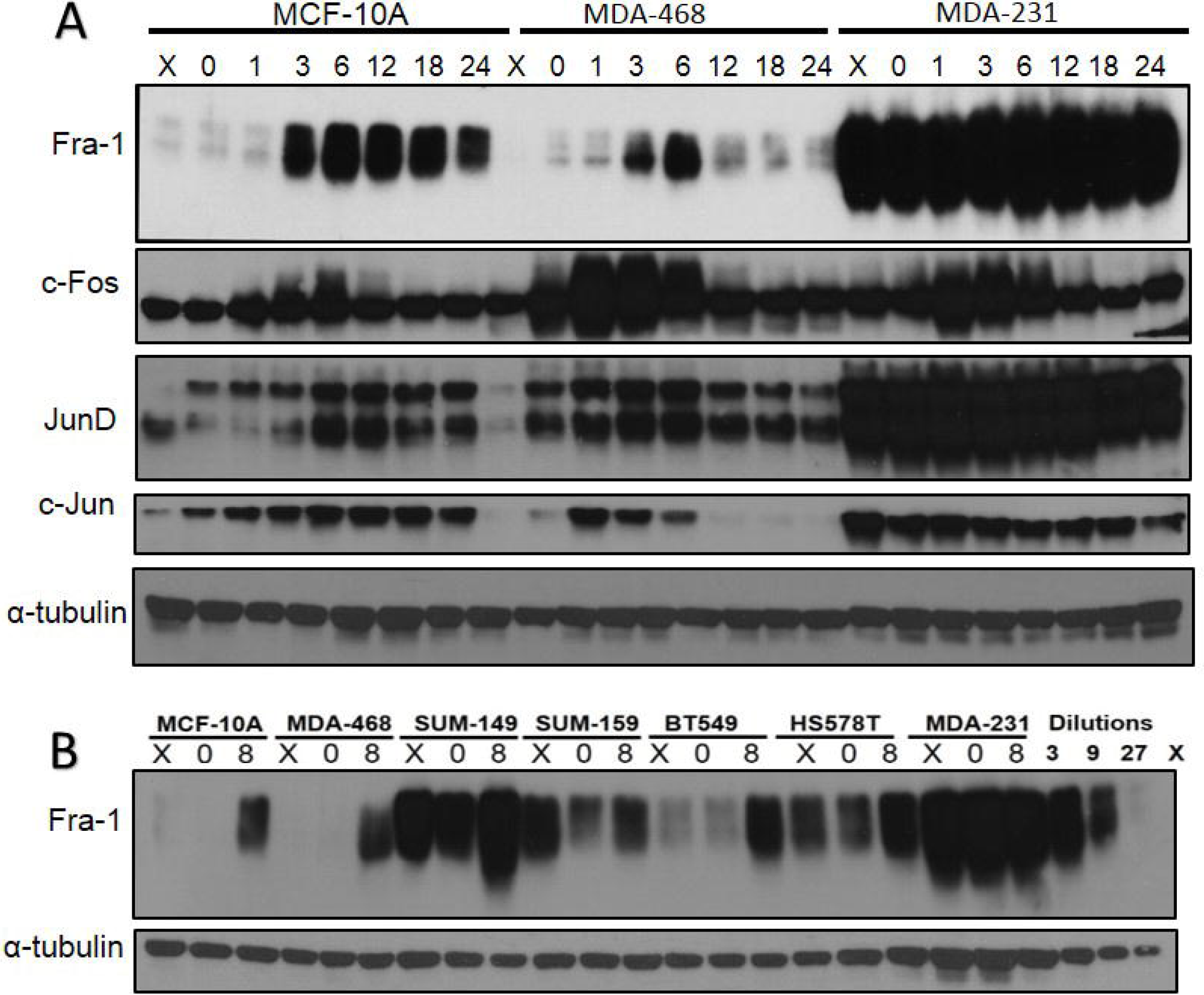
Analysis of Fos and Jun family members in non-tumorigenic and tumorigenic breast epithelial cell lines. (A) MCF10A, MDA-MB-468, and MDA-MB-231cells were subjected to serum starvation, then stimulated with serum for the times indicated in the figure (h). Cells were harvested and protein levels were detected using western blot. X = exponentially growing cells. The figure is representative of more than three independent experiments. (B) Fra-1 protein levels were analyzed in a panel of TNBC cell lines. X = exponential growth 0 = serum starvation for 48 hours. 8 = 8 hours of serum stimulation.

Additionally we sought to understand the distribution and dimerization pattern of different AP-1 members in MDA-MB-231 cells. So, we used nuclear fractionation to study Fra-1, c-Jun and JunD levels in the nucleus and cytoplasm in MDA-MB-231 cells. Our results showed that Fra-1, c-Jun and to less extent JunD are present both in the nucleus and cytoplasm of the MDA-MB-231 cells regardless of the cell cycle stage. (Figure S1). Then using co-immunoprecipitation (Co-IP) we found that both c-Jun and Jun-D dimerizes with Fra-1 in the nucleus. However in the cytoplasm, only c-Jun dimerizes with Fra-1 and to much lower extent than in the nucleus (Figure S1).

To examine if the high level of AP-1 members during serum starvation occurs in other TNBC cell lines, we tested Fra-1 level in a panel of TNBC cell lines compared to the non-tumorigenic MCF10A following serum starvation and release. The choice of Fra-1 was because it is the most studied member in breast cancer cell lines. All the examined cell lines reflected one of two patterns of Fra-1 expression (Figure 1B), and were categorized in two groups. The first group exhibit low levels of Fra-1 expression during serum starvation but induced by serum treatment. It includes BT549, Sum-159, MCF10A and MDA-MB-468. In the second group, which includes MDA-MB-231, SUM149, and HS578T, Fra-1 was expressed levels of Fra-1 during serum starvation or treatment.

### Fra-1 expression in cell lines of colon, prostate, lung and melanoma origin

To determine whether Fra-1 is highly expressed during serum starvation in cancer cell types other than breast cancer, Fra-1 levels were examined in colon, prostate, melanoma, and lung cancer cells following serum starvation and release. All the examined cell lines reflected one of two patterns of Fra-1 expression similar to those were detected in breast cancer (Figure 2). An exception was the lung cell line A549 that expresses low Fra-1 under both conditions. The colon cancer cells SW620 and SW480 belonged to the second group and exhibit more metastatic ability than the first group (Caco2 and HT29) based on results reported in previous studies [20, 21]. For cell lines of prostate origin, the first group is represented by the DU145 cell line that is less metastatic than PC-3 [22, 23] that represents the second group. For melanoma cells, the only information currently available is that Mel-147 cells has more migratory ability than Mel-19 cells [24]. These findings suggest that high Fra-1 expression in the absence of serum is predictive of the behavior of cancer cells across several types of cancer.

**Figure 2:**
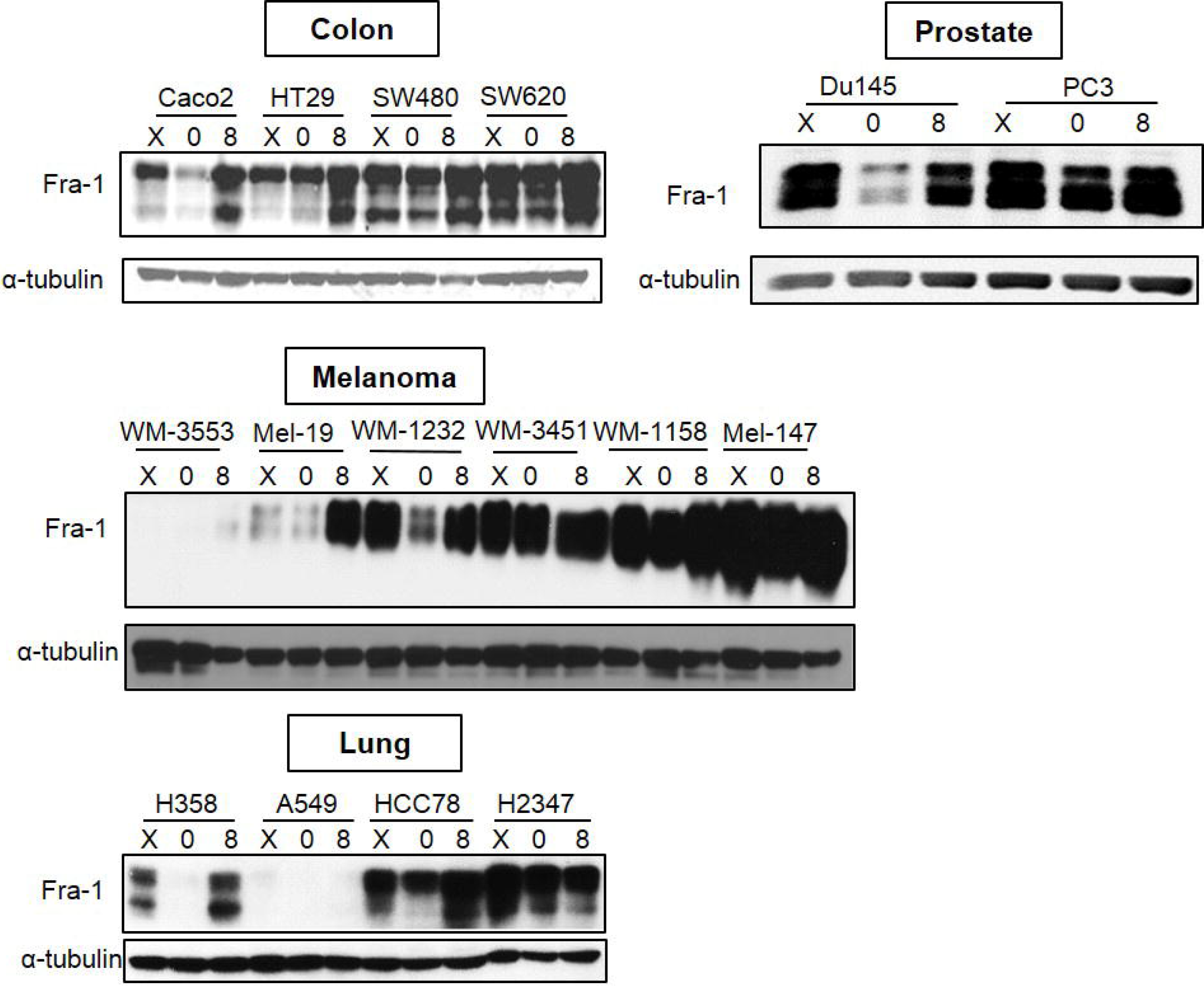
Analysis of Fra-1 expression in cancer cell lines of different origins. Colon, lung, prostate, and melanoma cancer cell lines were analyzed for Fra-1 protein by western blotting. X = exponential growth in complete medium. 0 = serum starvation for 48 hours. 8 = 8 hours of serum stimulation.

### High expression of Fra-1 correlates with cancer cell proliferation and migration in the absence of serum

To determine whether the differences in Fra-1 expression correlate with the ability of cells to cycle in the absence of serum, MCF-10A, MDA-MB-468 and MDA-MB-231 cells were serum starved, treated with nocodazole (NCD) for an additional 30 hours, and analyzed for cell cycle position by flow cytometry. Nocodazole blocks cells in mitosis, leading to an increase in the G2/M peak in flow cytometry. If cells are arrested by serum starvation, they will remain in G0/G1 during the nocodazole treatment. If they progress through the cell cycle despite being in serum starvati0n, they will be arrested at mitosis by nocodazole, and an increase in the G2/M peak will be observed. As shown in Figure 3A, serum starvation of MCF10A cells results in an efficient G0/G1 arrest. MDA-MB-468 and MDA-MB-231 cells also showed an increase in the percentage of G0/G1 cells in response to serum starvation, although to a lesser extent than MCF10A. Few if any MCF10A cells progressed to mitosis during the nocodazole treatment, confirming the strength of the G0/G1 arrest In contrast, MDA-MB-468 and MDA-MB-231 cells show continued advancement towards “S” phase and “G2/M” in the presence of nocodazole. However, the progression is significantly more evident in MDA-MB-231, where fewer cells remain in G0/G1 after starvation and nocodazole treatment. These results indicate a correlation between the high level of Fra-1 during serum starvation and the ability of these cells to progress through the cell cycle.

**Figure 3:**
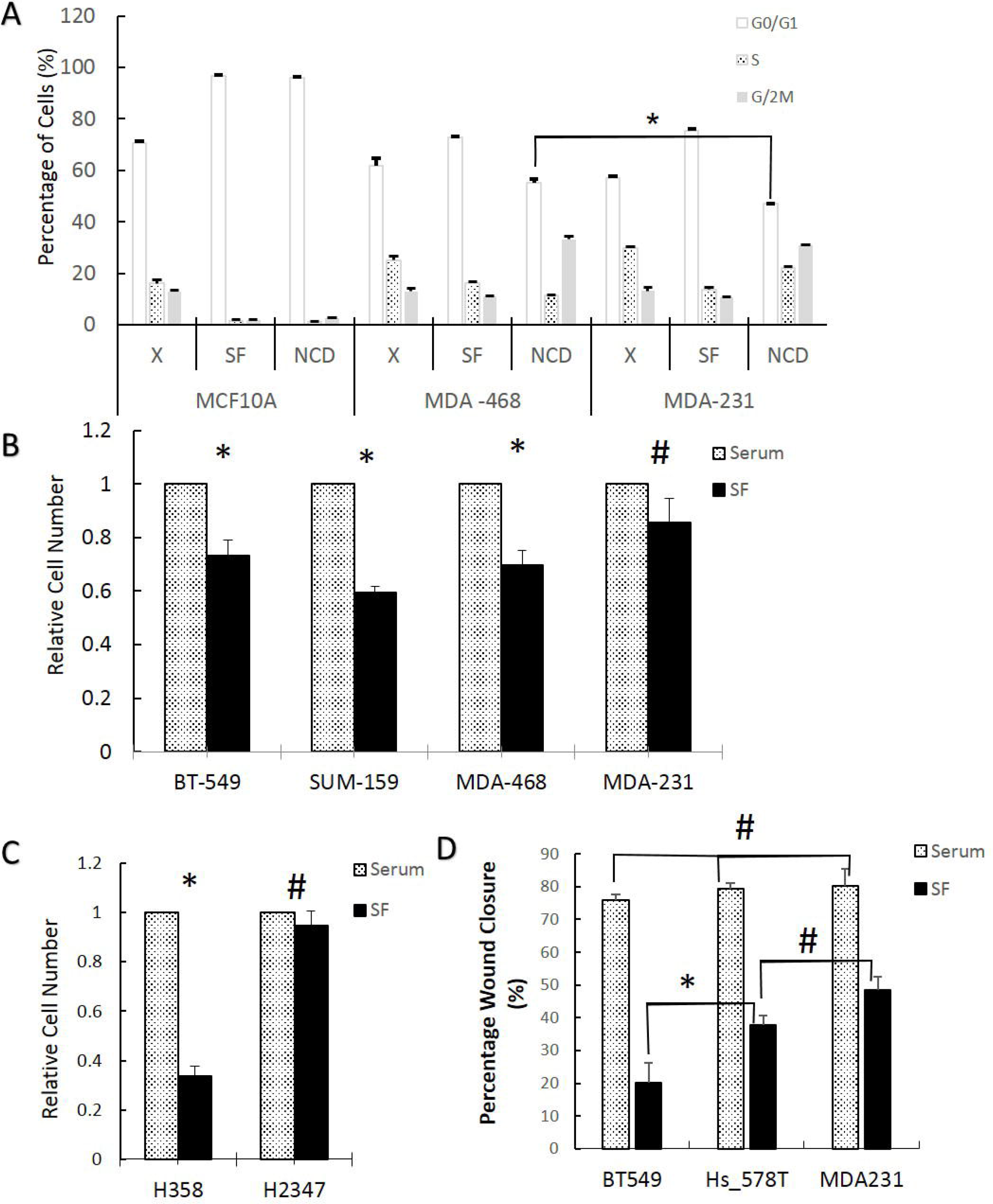
Fra-1 expression during serum starvation correlates with the ability of cells to proliferate and migrate in the absence of growth factors. (A): Cells were collected either during exponential growth (X), after 48 hours in serum free medium (SF), or following serum starvation plus treatment with nocodazole (250ng/ml) for 30 hours (NCD). Cells were stained with propidium iodide and analyzed for cell cycle distribution by flow cytometry. Data shown is the average of one experiment conducted in triplicate. (B and C): Cell growth in the presence or absence (SF) of serum was determined using manual cell counting as described in Materials and methods in breast (C) and lung (D) cancer cell lines. (E) Wound healing assays were performed in the presence and absence of serum in breast cancer cell lines. *= significant (p<0.05) using Student’s t test, # = non-significant. All numbers are the average of 3 independent experiments.

To strengthen the correlation between high expression of AP-1 family members and the ability of these cells to proliferate and migrate during serum starvation, we assayed the ability of several breast and lung cancer cell lines to proliferate and migrate in the presence or absence of serum. MDA-MB-231 increased in number to the same extent in the presence or absence of serum while in BT549, SUM159, and MDA-MB-468 cell proliferation was serum dependent (Figure 3B). Lung cancer H2378 cells grew equally in the presence or absence of serum while the H358 cells that show low Fra-1 grew only in presence of serum (Figure 3C). Finally, 3 different breast cancer cell lines were used to assess cell migration using a wound-healing assay in the presence and absence of serum. There was no significant difference in the rate of migration of the three cell lines in presence of serum. In contrast, when assayed in the absence of serum, cells that have high Fra-1 during serum starvation (HS578T and MDA-MB-231) had a significantly higher rate of migration than low Fra-1 BT549 (Figure 3D). These results show a correlation between the presence of Fra-1 and the ability of cancer cells to proliferate and migrate in absence of serum.

### Cancer cell proliferation and migration in the absence of serum is dependent on AP-1 activity and Fra-1 expression

The results above established a correlation between Fra-1 expression during serum starvation and the ability of cells to proliferate and migrate in these conditions. To determine if AP-1 activity is required for these phenotypes, we inhibited its activity by infecting MDA-MB-231 cells with a retrovirus vector that expresses a dominant negative form of c-Fos (A-Fos) under the control of a doxycycline (Dox) inducible promoter. First, we confirmed the expression of A-Fos in MDA-MB-231/Flag-AFos cells after induction with doxycycline using western blotting (Figure 4A). This blot also showed a reduction of Fra-1 after induction of A-Fos. Next, we asked whether the Fra-1 reduction due to A-Fos occurred in the cytoplasm and nucleus in presence or absence of serum by immunofluorescence staining. As shown in Figure S3, Fra-1 is both nuclear and cytoplasmic in MDA-MB-231 cells either in presence or absence of serum. After expression of A-Fos, total Fra-1 was reduced and it was located mostly in the cytoplasm.

**Figure 4:**
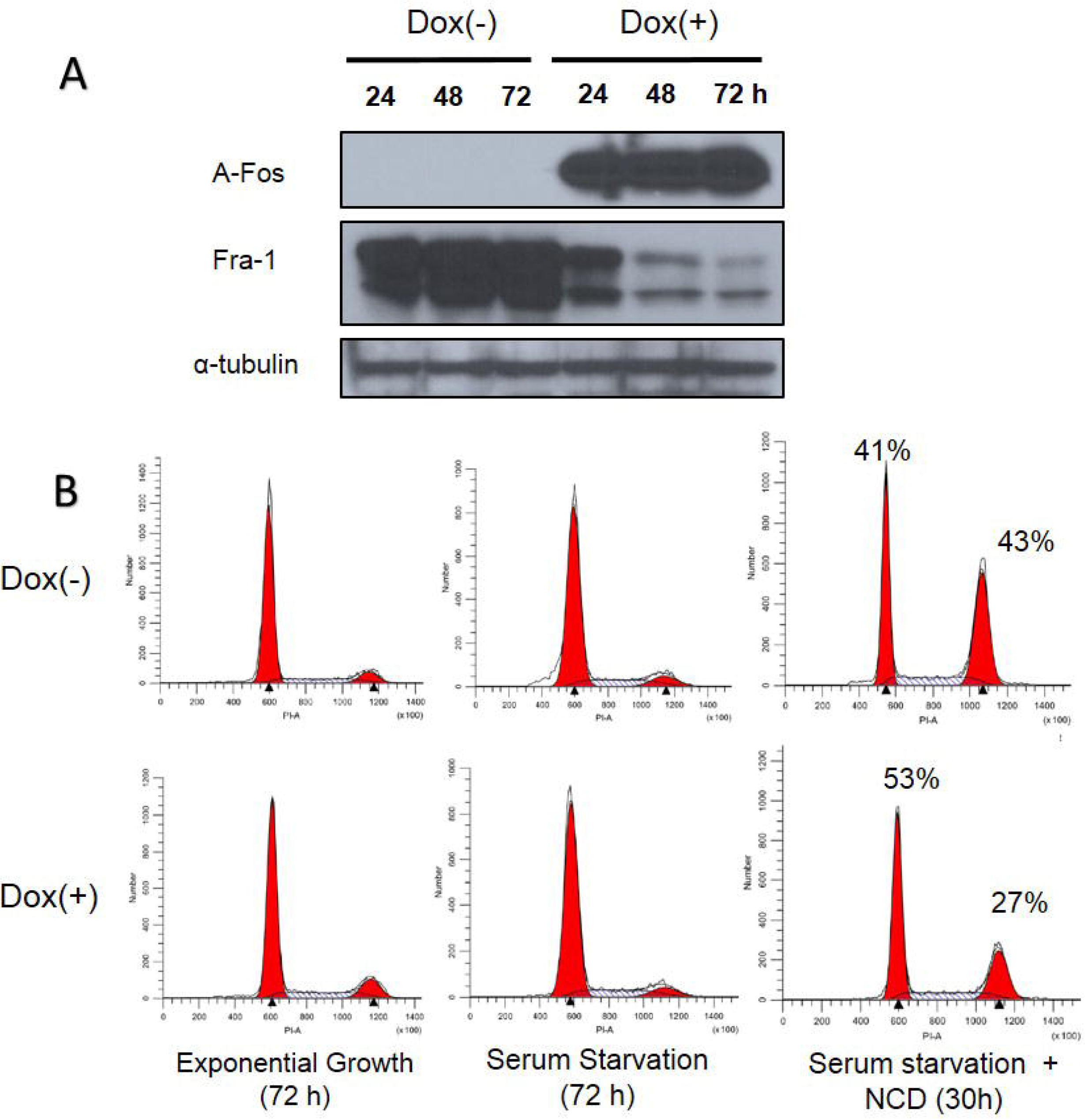
Inhibition of AP-1 activity in MDA-MB-231 cells reduces Fra-1 expression and suppresses the ability of the cells to progress through the cell cycle in the absence of serum. MDA-MB-231 cells were infected with a retroviral vector encoding a doxycycline inducible Flag-AFos gene as described in material and methods. Transfected cells were divided into two groups; non-induced Dox (−) or induced Dox (+). (A) A-Fos and Fra-1 protein levels were examined by western blotting. The figure is representative of 3 independent experiments (B) Cells were treated +/− Dox then subjected to serum starvation and nocodazole treatment as described in Figure 3. The data shown is representative of three independent experiments.

After A-Fos expression was confirmed, we examined whether AP-1 activity is required for cell cycle progression under serum starvation in MDA-MB-231 cells. As shown in Figure 4B, expression of A-Fos in MDA-MB-231 cells inhibited their progression through the cell cycle under serum starvation. This can be seen from the increase of cells arrested in G0/G1 and the decrease in accumulation of cells in G2/M when nocodazole is added. Next, we examined if the activity of AP-1 is required for cells to proliferate in absence of serum. MDA-MB-231/Flag-AFos cells were cultured in presence or absence of serum then Doxycycline was added in one group (Dox (+)) and the other group was used as a control (Dox (−)) (Figure 5A). Expression of A-Fos significantly reduced the number of cells in the presence of serum. However, in the absence of serum cell number was reduced but the reduction was not statistically significant. The cells were re-plated at the same initial density and incubated for another 72 hours (Second passage). This time the reduction was significant both in presence and absence of serum (not shown). When the cells were re-plated for the third time (Third passage), the effect was even stronger, after which the cells went into crisis with only very few cells surviving. When the surviving cells were allowed to grow they failed to express A-Fos (Data not shown).

**Figure 5:**
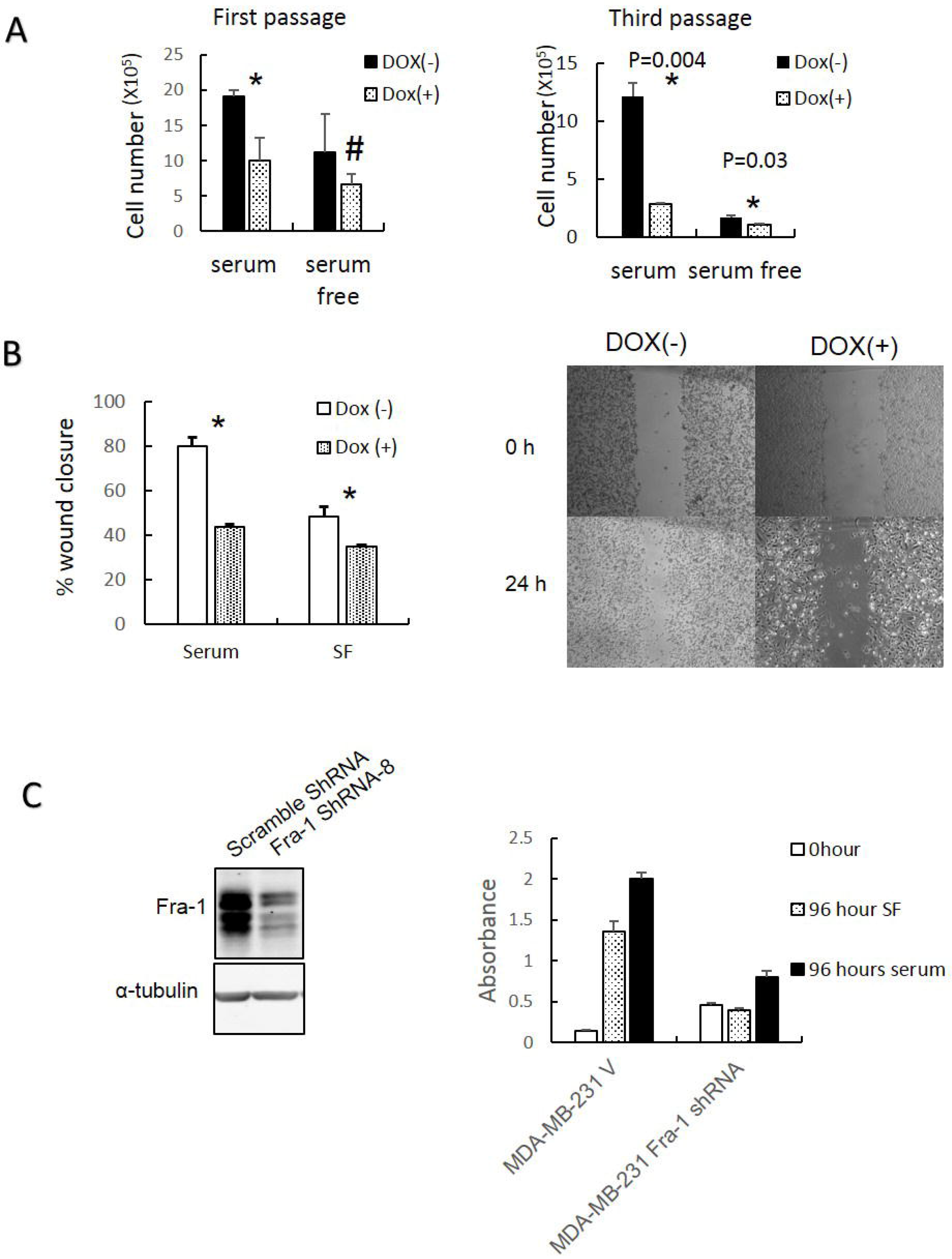
Inhibition of AP-1 activity and Fra-1 expression reduces MDA-MB-231 cell growth and migration. (A) The effects of A-Fos expression on the ability of the MDA-MB-231 cells to grow in the presence and absence of serum were analyzed for several passages. Cells were plated at density of 3×10^5^ with and without Dox and/or serum. After 72 hours cells were counted and re-plated at the same density, and this process was repeated for 3 passages. (B) The effect of A-Fos on the migration of MDA-MB-231 cells in the presence and absence of serum using the wound healing assay *= significant (p<0.05), the supplementary image represents migration in presence of serum. (C) Cells were infected with a retroviral vector encoding a scrambled sequence (Scramble shRNA) or Fra-1 shRNA (Fra-1 shRNA-8). Protein lysates were analyzed by western blot to detect Fra-1 expression (left). Cell proliferation was compared in absence (96 hour SF) and presence (96 hours of serum) of serum using CCK8 cell counting kit. The (0 hour) represents the absorbance at time of plating (Right).

Additionally, we tested the effect of A-Fos on the ability of MDA-MB-231 to migrate in presence and absence of serum using a wound healing migration assay. A-Fos was able to suppress cell migration both in presence and absence of serum (Figure 5B). These results indicate that AP-1 activity is required for cell proliferation and migration both in the presence and absence of serum.

To examine if Fra-1 is specifically required for promoting proliferation in MDA-MB-231 in absence of serum, we established stable derivatives of MDA-MB-231 expressing Fra-1 shRNA or a scrambled shRNA control, and examined the ability of these cells to proliferate in the presence or absence of serum. As shown in Figure 5C, growth of cells expressing Fra-1 shRNA was greatly decreased relative to controls, indicating that Fra-1 is important for proliferation.

### AP-1 dependent soluble factors from MDA-MB-231 cells induce Fra-1 expression in MCF10A and MDA-MB-468 and enhance their migration

The mechanism(s) by which Fra-1/AP-1 promotes proliferation and migration in the absence of serum are not well understood. We hypothesized that MDA-MB-231 cells produce soluble factors that might have a role in such a mechanism. To test this hypothesis, MDA-MB-231 cells were co-cultured with MCF10A or MDA-MB-468 cells on opposite sides of a trans-well insert. As shown in Figure 6A and 6C, this led to increased Fra-1 levels in both MCF10A and MDA-MB-468 cells. The effect of conditioned medium (CM) from serum deprived MDA-MB-231 cells on MCF10A and MDA-MB-468 cells was also examined, and resulted in increased Fra-1 expression in both cell lines (Figure 6B, C). To determine if AP-1 activity is required for this effect, the Dox-inducible MDA-MB-231/A-Fos cell line was utilized. CM from Dox-induced MDA-MB-231/A-Fos cells lost the ability to increase Fra-1 levels in MCF10A cells compared to the non-induced cells (Figure 6 D). Similar results were obtained using MDA-MB-468 cells (data not shown). Because the MEK/ERK pathway is the canonical pathway that regulates AP-1 expression, we explored its role in the induction of Fra-1 in MCF10A cells by conditioned medium. Two different MEK inhibitors were used; U0126 and PD-98059. Both MEK inhibitors decreased the effect of conditioned medium on the level of Fra-1 in MCF10A cells (Figure 6E). These results indicate that AP-1 dependent soluble factors from MDA-MB-231 cells induce Fra-1 expression in MCF10A and MDA-MB-468 cells, likely through activation of the MEK/ERK pathway.

**Figure 6:**
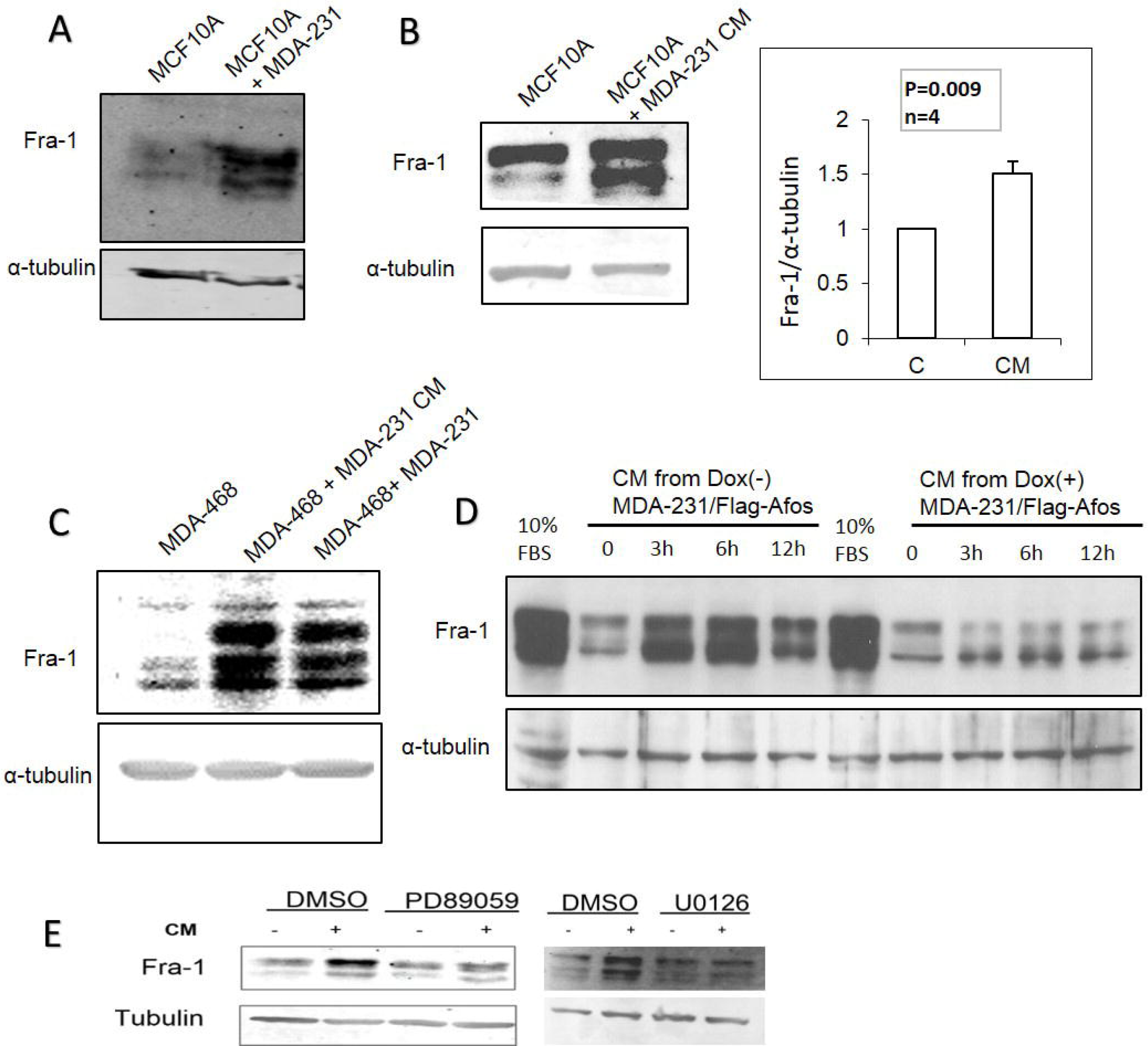
AP-1 dependent soluble factors from MDA-MB-231 cells induce Fra-1 expression in MCF10A and MDA-MB-468 cells. (A) Using a transwell system MCF10A cells were cultured in the upper chamber with MDA-MB-231 cells or serum free medium in the lower chamber, then the levels of Fra-1 protein in MCF10A cells were examined. (B) Conditioned medium from serum starved MDA-MB-231 cells was added to serum starved MCF10A cells for 6 hours then Fra-1 protein levels were examined. Blots were quantified using Image J software. (C) MDA-MB-468 cells were incubated with CM or cocultured with MDA-MB-231 cells in a transwell chamber as described in materials and methods, then Fra-1 expression was examined. (D) CM from MDA-MB-231/Flag-AFos cells incubated in the presence (Dox(+)) and absence (Dox(−)) of doxycycline was added to MCF10A cells, and Fra-1 protein levels were examined. The results shown are representative of 3 experiments. (E) MCF10A cells were incubated with MDA-MB-231 CM with and without PD98059 and U0126 (MEK inhibitors). The cells were harvested after 6 hours and the level of Fra-1 was detected by western blotting.

AP-1 controls the expression of many genes that enhance cell migration and metastasis [25]. We therefore tested whether co-culture with MDA-MB-231 and/or MDA-MB-231 CM treatment of MCF10A cells increases their ability to migrate in transwell assays. As shown in Figure 7A and B, migration of MCF10A cells increased when MDA-MB-231 cells were in the lower chamber, and treatment with conditioned medium from MDA-MB-231 cells led to a similar result. The CM experiment was conducted in two different scenarios. Conditioned medium or serum free medium as a control, was added in the upper chamber and cells were allowed to migrate towards 10% FBS in the lower chamber. Alternatively, the cells were grown in serum free medium in the upper chamber then allowed to migrate toward serum free medium versus conditioned medium in the lower chamber. In both cases, the number of migrating cells significantly increased with CM as compared to the control. Similar results were obtained with MDA-MB-468 cells when they were co-cultured with un-induced MDA-MB-231/A-Fos cells, and the effect was blocked by A-Fos induction (Figure 7C). The effect of MDA-MB-231 CM on MCF10A cell migration was also demonstrated in a wound healing assay (Figure 7D). To determine if Fra-1 expression in MCF10A cells is required for the response to CM, Fra-1 shRNAs were expressed in MCF10A cells. Two independent shRNAs showed knockdown of Fra-1, and were used to produce stable cell lines. Knockdown of Fra-1 in MCF10A significantly reduced the migratory response to MDA-MB-231 CM (Figure 7E). Thus, Fra-1 is required for both the production of and the response to the soluble factors in MDA-MB-231 CM.

**Figure 7.**
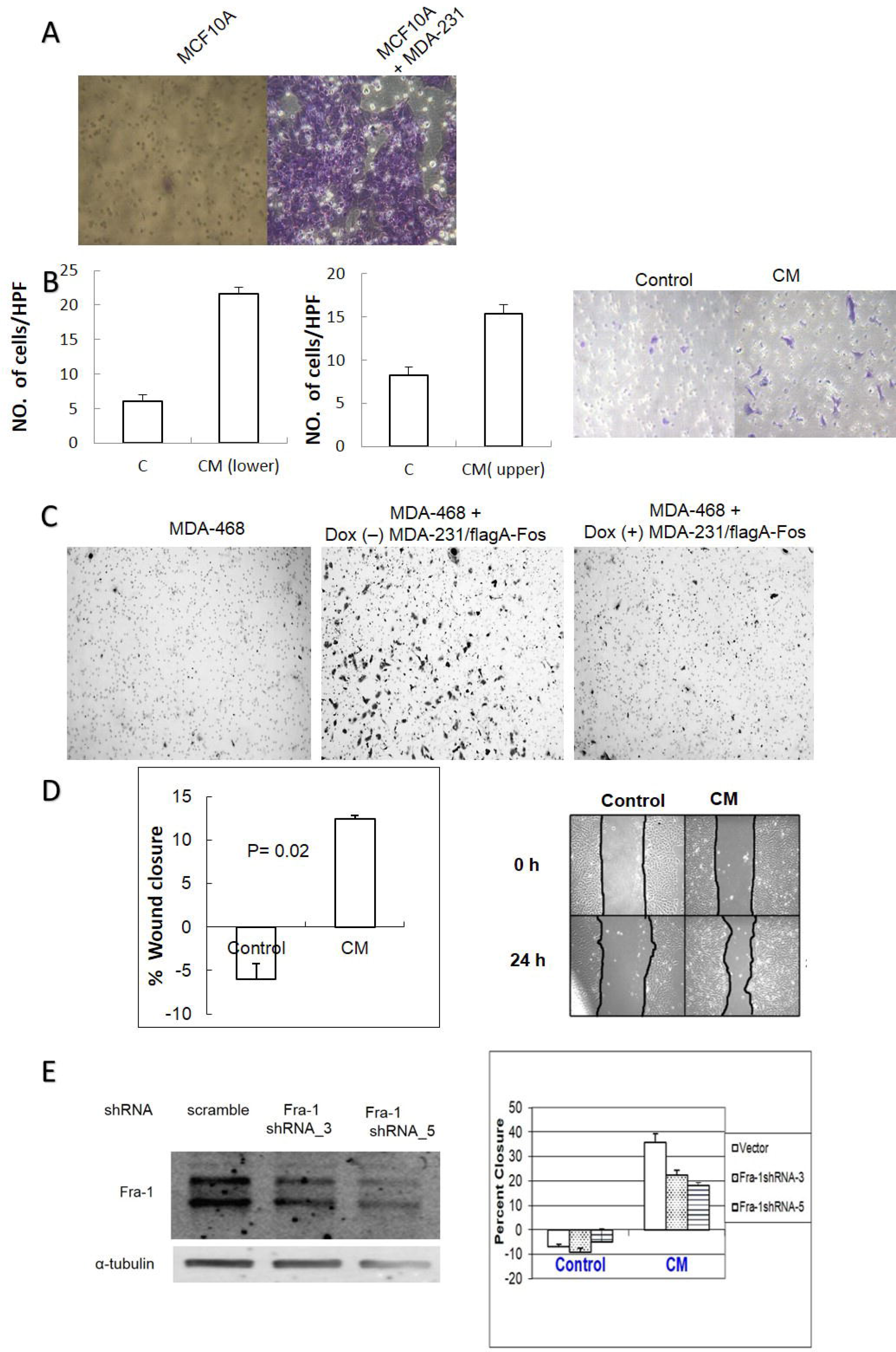
AP-1 dependent soluble factors from MDA-MB-231 cells increase migration of MCF10A and MDA-MB-468 cells in an AP-1 dependent manner. (A) MCF10A cells were cultured on an 8μm pore transwell insert with or without MDA-MB-231 cells in the lower chamber and migrated MCF10A cells on the lower side of the membrane were fixed and stained as described in materials and methods. (B) MCF10A cells were cultured in the upper chamber of transwell inserts in MDA-MB-231 CM versus serum free medium (left graph), or MCF10A cells were cultured in the upper chamber in serum free medium with the lower chamber filled with either CM or serum free medium (right graph). Cells were counted per HPF. from 3 independent experiments and a representative picture of the migrated cells is shown. (C) MDA-MB-468 cells were co-cultured with MDA-MB-231/Flag-AFos cells in a transwell system in the presence (Dox(+)) and absence (Dox(−)) of doxycycline. (D) A wound healing migration assay was used to measure the effect of CM on MCF10A cell migration. The graph shows percent closure after 24 hours. (E) MCF10A cells were infected with scrambled shRNA virus (vector) or two different Fra-1 shRNA viruses and Fra-1 levels were compared (Left). A wound healing assay was carried out to measure the effect of conditioned medium on MCF10A migration (Right).

## DISCUSSION

AP-1 family members have been associated with several oncogenic properties of cancer cells from different tissue origins [15, 26–38]. Amongst them, Fra-1 and c-Jun were the most clearly associated with tumor progression in breast [16, 17], colon [39], prostate [40], lung [41] cancers and melanoma[42]. Fra-1 and c-Jun mediated increased proliferation, migration, invasion and epithelial mesenchymal transition (EMT) [32] and inhibited apoptosis [34] of these cancers.

The main objective of the current work was to decipher the role of AP-1 in maintaining cancer cell growth and migration in absence of mitogenic growth factor signaling. Our findings indicate that AP-1 activity and Fra-1 expression contribute to the ability of cancer cells to grow and migrate in the absence of growth factors; two fundamental hallmarks of cancer [19]. In addition, we suggest that the ability to proliferate and migrate in the absence of serum is mediated via self-secreted factors, and that both the synthesis of and response to these factors is dependent on AP-1 signaling.

To the best of our knowledge, the current study is the first to compare AP-1 regulation during reentry into the cell cycle in mammary epithelial cells and breast cancer cell lines. All previous studies were conducted in rat fibroblasts and RAS transformed cell lines [9, 10, 43–45]. Our results showed that the pattern of expression of AP-1 members in MCF10A and MDA-MB-468 is similar to previous results in rat fibroblasts. The most prominent feature in our comparison was the persistently high level of Fra-1 and c-Jun in MDA-MB-231 cells in the complete absence of serum that was maintained at all stages of the cell cycle. Similar to our results, RAS transformed NIH3T3 cells showed higher level of Fra-1 and c-Jun that persisted even in serum starved cells [46].

Our results suggest a novel role for AP-1 in maintaining the ability of breast cancer cells to continue to grow and migrate in absence of serum. Also our results extend this role to other types of cancer like colon, prostate, lung cancers and melanoma. Previous work showed that when Fra-1, Fra-2 or c-Jun are overexpressed in non-tumorigenic or non-metastatic cell lines, their effect was masked if the cells are grown in presence of serum [17, 46]. Also, Bakiri and colleagues [47] found that when the tethered c-Jun~Fra-2 dimer was overexpressed in NIH3T3 cells, these cells were able to proliferate in absence of serum. We conclude that, in presence of serum or growth factors, AP-1 family members are high and give non-invasive cells the ability to proliferate and migrate that is comparable to that of the cancer cells. However, invasive cells that express high levels of AP1 family members in the absence of serum are able to proliferate and migrate even when they are deprived of growth factors. Constitutive induction of transcription factors that mediate such functions is one of the mechanisms utilized by cancer cells to maintain their functions in absence of growth factors [19]. In this work we established the role of AP-1 in this context.

Another major finding of our study is the ability of MDA-MB-231 to secrete AP-1 inducing factors. We propose that these factors are responsible for paracrine or autocrine loops, and represent a possible mechanism to keep cells proliferating and/or migrating in absence of growth factors. A paracrine role was detected by the ability of CM from MDA-MB-231 cells or co-culture with MDA-MB-231 to induce Fra-1 and enhance migration in MCF10A or MDA-MB-468 cells. Secretion of these factors is AP-1 dependent, as demonstrated by the ability of A-Fos to inhibit its action. The ability of these factors to induce migration of MCF10A cells was also dependent on Fra-1 expression in the recipient cells. This indicates that Fra-1/AP-1 could mediate the action of soluble factors that are secreted by the MDA-MB-231 cells. However, the direct proof that these factors function in an autocrine manner to induce MDA-MB-231 cell proliferation/migration is yet to be obtained.

Previous studies showed the role of soluble factors in the aggressiveness of cancer cell lines and suggested a role for different pathways. For instance, Lieblein and colleagues [48] suggested a role for STAT3 in mediating such a loop. Our study provided evidence for a central role of AP-1 as a mediator of such a paracrine loop. Similarly, previous studies proposed a role of Fra-1 through EGF in lung cancer [49] or TGF in colorectal cancer [50]. Since there is evidence that AP-1 and STAT3 act synergistically to boost aggressiveness of cancer cells [51], AP-1 and STAT3 may cooperate to maintain such a loop.

Most studies of paracrine loops focus on the effect of cancer cells on adjacent stromal and immune cells [52, 53]. Minimal attention has been given to the effect of cancer cells on adjacent normal parenchymal cells and its role in enhancing cancer metastasis. This study is the first to show the ability of soluble factors from cancer cells to induce Fra-1 in non-tumorigenic mammary epithelial cells. This points to a role of cancer cells in inducing AP-1 factors in neighboring non-cancerous functional (parenchymal) cells which may enhance cancer metastatic potential by different mechanisms. For example, since Fra-1 promotes EMT [32], the increase in its expression in normal mammary epithelial cells surrounding a tumor may compromise the epithelial barrier and facilitate metastasis. Another possible mechanism is through enhancing the secretome of these neighboring cells by a paracrine action leading to reciprocal activation of cancer cells. AP-1 is known to induce the secretion of several cytokines such as EGF [54], TGFB [55], osteopontin, IL-6, IL-8, VEGF and extracellular enzymes [56] that promote cancer metastasis. The paracrine induction of AP-1 in neighboring cells is expected to enhance the secretion of these metastasis promoting factors from these cells. This will further support the metastatic potential of the tumor. In addition, paracrine signaling by tumor cells may contribute to their ability to colonize specific tissue types during the metastatic process.

Our results and previous studies suggest a universal function of Fra-1 in metastatic cancers, and that it might serve as a target for universal cancer treatment. Despite the current focus on individualized therapy, scientists are still hoping to find a universal treatment for cancer [57]. Since transcription factors are not readily druggable, an alternative strategy that has been proposed is to look for an upstream regulator or downstream effector of AP-1 [58] [59]. Characterizing an autocrine loop that controls AP-1 function may identify targets that fulfill both roles. In addition, recent studies describe approaches to target transcription factors including oligo deoxy-nucleotides [60] and deoxyribozymes [61]. Here we demonstrated the ability of A-Fos to suppress both growth and migration of the highly aggressive MDA-MB-231 cell line. We also detected the ability of A-Fos to inhibit anchorage independent growth of MDA-MB-231 cells (Data not shown). Based on our findings, we propose that an alternative strategy would be the use of A-Fos as a potential gene therapy tool.

In summary, our results demonstrate a fundamental role of AP-1 family members in cancer cells. First, they identify the role for AP-1 in maintaining cancer cell proliferation and migration in absence of growth factors. Additionally, this work highlights a central role of AP-1 in the secretion of and response to autocrine/paracrine factor(s) that play an important role in enhancing cancer cell proliferation and metastasis. This pivotal role of AP-1 empathizes its position as an important target for cancer therapy.

## MATERIALS AND METHODS

### Cell lines

The breast cancer cell lines MDA-MB-231 (a gift from Dr. Kathleen Gallo, Michigan State University (MSU)), and MDA-MB-468 (a gift from Dr. Chengfeng Yang, MSU) were maintained in DMEM medium with 10% FBS. The BT549, SUM149 and SUM 159 were gifts from Dr. Chengfeng Yang (MSU). The MCF10A cell line (a gift from Dr. Susan E. Conrad, MSU) was maintained in DMEM/F12 medium supplemented with 5% horse serum (HS) (Atlanta Biologicals), 20 ng/ml Epidermal Growth Factor (EGF) (Sigma, St. Louis, MO), 100 ng/ml Cholera Toxin (CT) (Sigma), 10 μg/ml Insulin (INS) (Sigma St. Louis, MO), 500 ng/ml hydrocortisone (HC) (Sigma), and 2.5 mM l-glutamine (Mediatech St. Louis, MO). The prostate cancer cell lines DU145 and PC3 cell lines were a gift from Dr. Peter C. Hollenhorst (Indiana University). The former was grown on Eagle’s minimum essential medium (EMEM) supplemented with 10% FBS while the later was grown on F12K medium (Life Technologies, 21127-022) with 10% FBS. All colon cancer cell lines (Caco2, HT29, SW480 and SW620) were obtained as a gift from Dr. Marc D Basson (University of North Dakota). Caco2 cells were grown in DMEM medium with 20% FBS while SW620, SW480 and HT29 were grown in RPMI medium with 10% FBS. Lung cancer cell lines (H2347, HCC78, A549, and H358) were obtained as a gift from Dr. Dipankar Ray (University of Michigan). All of them were grown in RPMI medium with 10% FBS. HFK293GPG cells (a gift from Dr. Kathleen Gallo, MSU) were grown in DMEM medium with 10% FBS. All cell lines were grown in 5% CO2 at 37 °C in a humidified incubator.

### Antibodies

The following antibodies were purchased from Santa Cruz Biotechnology (Santa Cruz, CA, USA): Anti-Fra-1 (sc-183) (sc-605x) (sc-28310), anti-c-Fos (sc-52), anti-c-Jun (sc-44) (sc-74543), anti-Jun D (sc-74) (sc-271938), anti-α-Tubulin (sc-8035), normal rabbit IgG (sc-2027), and mouse-IgG_k_ BP-HRP (sc-516102). Goat anti-rabbit IgG HRP (ADI-SAB-300-J) was purchased from Enzo Life Science, Inc. (Farmingdale, NY, USA). ECL Sheep anti-Mouse IgG HRP (NA931V) was from Amersham Bioscience UK limited (Buckinghamshire, UK). IRDye^®^ 800CW Donkey anti-Rabbit IgG (926-32213), IRDye^®^ 680RD Donkey anti-Mouse IgG (926-68072) were from Li-COR Bioscience (Lincoln, NE, USA). Goat anti-Rabbit IgG conjugated with Alexa Flour 488 was from Thermo Fisher Scientific, Inc (Invitrogen) (Waltham, MA, USA).

### Cell cycle re-entry

Exponentially growing MCF10A, MDA-MB-231, and MDA-MB-468 cells were washed twice with serum free medium and incubated with 0.05 % serum in DMEM for either 48 hours (MDA-MB-231 and MDA-MB-468) or 24 hours (MCF10A). Following serum starvation, one group of cells was harvested. For the rest of the cells the medium was replaced with medium containing 10% FBS and further cultured until harvested after 1,3,6,12,18 or 24 hours.

### Western blot

Whole cell protein lysates were prepared using lysis buffer (50 mM Tris-HCL pH 7.5, 150 mM NaCl, 1 mM EDTA, 2 mM EGTA, 30 mM NaF, 10 mM NaPP, 1% Triton X-100, 10% Glycerol, 0.5% Deoxycholate) supplemented with protease inhibitor cocktail, phosphatase inhibitors and SDS (0.01%). Quantification of total protein was assessed using Biorad DC protein assay kit (Biorad 500-0116) and 20 μg of total protein from each sample was separated on 12% SDS PAGE gel then transferred to PVDF membrane (BioRAD, 162-0177). Non-specific binding was blocked with PBS containing 5% dry milk for 1h. The membrane was probed for 1h at RT with PBS containing 0.1 Tween20, 0.05 % milk and the primary antibody, then with the appropriate secondary antibody conjugated with horseradish peroxidase or IRDye and developed by the chemiluminescence method or by fluorescence, respectively.

### Cell cycle analysis

Individual pellets of freshly trypsinized cells (1-2 ×10^6^ cells) were re-suspended in 400μl PBS/FBS mixture (1:1) and fixed by adding 1200 μl of 70% cold ethanol to achieve a 50% final ethanol concentration. The cells were stored on ice for at least 5 hours then washed twice in 5% heat inactivated (HI) calf serum in PBS (pH 7.2), and centrifuged at 1400 RPM for 5 min at room temperature. The cells were resuspended in 1 ml of PBS (pH 7.2) containing 50μg/ml propidium iodide (Sigma. Cat. No. P4170), 5% HI calf serum, and 1mg/ml of RNase A, then incubated for 15 min at 37°C. Cell cycle analysis was performed using flow cytometry (LSRII BD bioscience) and Verity software House Modfit 4.1.

### Nocadazole block

Cells were plated in 10 cm dishes in three groups; exponential, serum starved, and nocadazole treated (Sigma-Aldrich, St. Louis, MO). Exponential cells were harvested first, and the other groups were serum starved for 48 hours. The he serum starved group was then harvested, and the remaining group (NCD) were treated with nocadazole (final concentration 250 ng/ml) for 30 h. All harvested cells were analyzed for cell cycle using flow cytometry.

### Cell proliferation assay by cell count

Cells were plated at a density of 3×10^5^ cells/ well in 6 well plates with DMEM containing 10%FBS. Once fully attached, the cells were washed twice with serum free medium and incubated with DMEM containing either 10% or 0.05% FBS. After 72 hours, cells were trypsinized and counted manually using a Bright Line hemocytometer (Reichert, Buffalo, NY).

### Wound healing migration assay

MCF10A cells were seeded in 6 well plates at a density of 3×10^5^ cells / well to achieve 95% confluence. Once attached, the cells were washed twice with serum free medium, and cultured in serum free medium for 24 hours. Vertical wounds were made in each well using P200 pipette tips. The wounded cells were washed once with PBS, and fed with either conditioned medium or serum free medium for 24 hours in the presence of mitomycin C (Sigma-Aldrich, St. Louis, MO). Cells were imaged using a Nikon Eclipse TS100 inverted microscope (Nikon). Random fields in each well were marked and imaged at the same location at 0 and 24 hours with a Ph1 ADL 10X/0.5 objective using a Cool snap Easy camera (Horiba Scientific) controlled by NS-Element D 3.1 Acquisition (Nikon) software. Images were analyzed and the area of the wound was calculated using ImageJ software. The percentage of wound closure was calculated as follows:

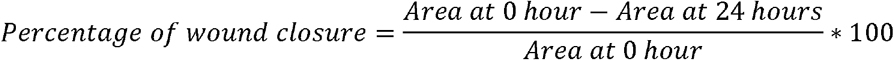

### Plasmids and Stable MDA-MB-231/Flag-AFos cell line

To generate inducible lentiviral vector pINDUCER20/Flag-AFos, the Flag-AFos fragment was excised as a 321bp Sal1-EcoR1 from pCMV-500 Flag-AFos (a gift from Dr. Richard Schwartz, MSU) and cloned into Sal1-EcoR1 cleaved pENTR1A (a gift from Dr. Brian Schaffhausen, Tufts Medical School) to generate pENTR1A-Flag-AFos. Flag-AFos was then recombined into pINDUCER20 (a gift from Dr. Brian Schaffhausen, Tufts Medical School) from pENTR1A-Flag-AFos using Gateway LR clonase II enzyme mix (Life Sciences). Lentiviral supernatants were generated by transient transfection of psPAX2, pMD2.G (a gift from Dr. Kathleen Gallo, MSU), and pINDUCER20-Flag-AFos into 293T cells according to the Invitrogen Lipofectamine 2000 transfection protocol. Medium containing the virus was harvested 48 h after transfection. Stable MDA-MB-231 cell lines expressing Flag-AFos were generated (MDA-MB-231/Flag-AFos) by lentiviral transduction in the presence of 8μg/ml polybrene followed by selection with G418 [62].

### Generation of Fra-1 shRNA stable knock down cells

MDA-MB-231 or MCF10A cells were transduced with retroviral particles expressing Fra-1 shRNA (LPE sh.F0SL1.1405 or LPE sh.FOSL1.1223) or scrambled control (LPE sh.Ren.713) (gifts from Mirimus Inc). To generate retroviruses, 293 GPG packaging cells were plated at a density of 1×10^6^ in 6 well plates. Twelve hours after removal of antibiotics, cells were transiently transfected with 1μg of retrovirus vector using Lipofectamine 2000 (Invitrogen). Medium was changed 24 hours following transfection. Cell supernatant was collected on days 4 through 7 after plating then filtered through 0.45 μm syringe filter, aliquoted, and frozen at −80 °C.

To infect the cells, they were plated overnight in 6 well plates at density of 5×10^4^ well. On the day of infection, 500μl of the virus containing medium was added to 500 μl of growth medium with 8 μg/ml of polybrene and the mixture was added to the cells for 4 to 5 hours hen 1 ml of growth medium was added. Twelve hours after the infection, cells were treated with 1μg/ml of puromycin to select for cells that stably expressed scrambled shRNA or Fra-1 shRNA. The decreased level of Fra-1 was confirmed by western blot.

### Cell proliferation assay by CCK8 cell counting reagent

Cells were plated in 3 groups in 96 well plates at a density of 1.5×10^4^ /well. The first group was used for measuring the initial reading at 0 hour. Cells of second and third groups were washed twice with serum free medium then 10% or 0.05% FBS medium was added. After the designated time period, the CCK-8 reagent (Dojindo Molecular Technologies, Inc. Cat. No.CK04-05) was added. After 1 hour, the absorbance was measured at 450 nm.

### Co-culture

Co-culture was conducted in 0.4 μm pore P.E.T membrane inserts (Falcon. Cat. No. 353090). The membranes were incubated in DMEM for initial equilibration for at least 1 hour. MDA-MB-231 were cultured on the membrane at a density of 2 ×10^5^ cells/well. After attachment to the membrane the medium was changed to 0.05% serum overnight. At the same time either MCF10A or MDA-MB-468 cells were plated at a density of 5 ×10^5^ cells/well in a 6 wells plate. When the cells reached 70 % confluence, 0.05% FBS medium was added to the cells. After 16 hours the trans-wells containing the MDA-MB-231 were transferred into the wells that contain the MCF10A or MDA-MB-468 cells, incubated for 24 hours then harvested in lysis buffer. Cell lysates were collected for Western blotting. One well of each cell type had no insert as a control.

### Preparation of conditioned medium

MDA-MB-231 cells were plated in 6 well plates at a density of 7.5 × 10^5^ cells / well and allowed to attach overnight. The cells were then washed twice with serum free medium and incubated for 24 hours in 1.5 ml DMEM with 0% FBS. The conditioned medium was harvested and used for subsequent culture assays.

### Transwell migration assay

Cell migration assay was carried out in Transwell chambers with 10.5 mm membrane diameter and 8 μm pores (Falcon. Cat. No. 353182). The membranes were pre-equilibrated in DMEM for 1 hour. MCF10A or MDA-MB-468 were cultured in DMEM with 2% serum for 24 h then 2×10^5^ cells were plated in each migration chamber. Once cells adhered to the membrane, the medium in the upper and lower chambers was changed to the appropriate medium. After 24 h, membranes were washed twice with cold PBS, cells in the upper surface of the membrane were removed with a cotton swab, and cells situated on the lower side of membrane were fixed for 20 minutes in PBS containing 3.7% formaldehyde. Cells were stained with 0.5% crystal violet in 10 % ethanol for 10 minutes then washed with tap water. The filters were photographed under 200X microscopic power and the number of cells was counted per high power field (HPF).

## ABBREVIATION

Activating Protein-1 (AP-1), Triple Negative Breast Cancer (TNBC), co-immunoprecipitation (Co-IP), nocodazole (NCD), a dominant negative form of Fos (A-Fos), doxycycline (Dox), serum free medium (SF), Condition Medium (CM), epithelial to mesenchymal transition (EMT), high power field (HPF).

## AUTHOR CONTRIBUTIONS

SAFI and AA: Planned and executed the experiments, analyzed and interpreted data, drafted the manuscript and did final corrections. EJ: Planned and executed the experiments, analyzed and interpreted data, and shared in manuscript revisions. NA: Planned and executed the experiments and analyzed and interpreted data. MF shared in study design and experimental planning, shared in data analysis and interpretation and contributed to the writing of the final version of the manuscript.

## ACKNOWLEDGMENT

We greatly acknowledge Prof Lakshmishankar Chaturvedi for his generous advice in experimental design and analysis of data. We would like to extend all the thanks to Prof. Suzan Conrad for her valuable discussions and revisions of this manuscript. We are also thankful to Dr. Jeanine Scott and Prof. Brian Schaffhausen for their valuable comments and corrections of the manuscript.

